# Extended pregnancy during matrotrophy in cockroaches increases progeny survival during dry periods

**DOI:** 10.64898/2026.08.26.745935

**Authors:** Gabrielle LeFevre, Jacob Hendershot, Sophie Shemas, John Cavanaugh, Luke Bernhardt, Matthew Korthauer, Ronja Frigard, Emily C. Jennings, Alexandra Jansen van Rensburg, Sinead English, Joshua B. Benoit

## Abstract

Animals reproduce across a spectrum from oviparity to viviparity. Internal gestation and live birth have inherent advantages despite their costs. However, comparative analyses of key drivers, such as resilience to dehydration, lack taxonomic breadth. In this study, we assessed whether live birth improves species’ ability to proliferate in environments with limited access to water. To do so, we used the viviparous cockroach, *Diploptera punctata*, as a model of viviparity to assess dehydration-induced damage during extended periods of water deprivation. During pregnancy, maternal survival did not decline during dry periods compared with non-pregnant individuals, suggesting that pregnancy has minimal impact on resistance to dehydration stress. When mothers are deprived of water for 2 weeks, they show an increase in osmolality after one week, while their embryos show no change in osmolality until after two weeks, at which point abortions occur. Periods of dehydration increased the duration of pregnancy, likely due to a slower production of *in utero* milk, but there was no change in the size of progeny. After birth, viviparous *D. punctata* newborns showed increased survival under dry conditions compared to two other ovoviviparous species, which are much smaller and dehydrate more quickly. The combined impact of increased dehydration resistance during pregnancy and during the first independent phase indicates a benefit of prolonged viviparity in *Diploptera punctata*. This study provides evidence that viviparity in animal systems could enable progeny survival during drought.

## Introduction

Viviparity, the reproductive strategy where females produce live young rather than laying eggs, has evolved more than a dozen times independently across diverse insect lineages, a striking example of convergent evolution (Fouks et al., 2023; Kalinka, 2015). This adaptation likely arose through gradual increases in egg retention within the female reproductive tract, followed by the evolution of maternal provisioning strategies that ensure embryonic survival (Gavrilov-Zimin, 2022; Kalinka, 2015; Ostrovsky et al., 2016). The spectrum of viviparity in insects ranges from ovoviviparity, in which eggs hatch inside the female with limited maternal input, to highly specialized forms of matrotrophy, in which the mother directly nourishes the developing offspring (Benoit et al., 2015; Ostrovsky et al., 2016). One of the best-studied examples of matrotrophy is adenotrophic viviparity in the tsetse fly (*Glossina* spp.), where a single larva develops within the uterus and is fed with specialized “milk” secretions until it is deposited as a fully grown larva ready to pupate (Attardo et al., 2019; Benoit et al., 2014a; Benoit et al., 2015). Some aphid species (e.g., *Acyrthosiphon pisum*) display viviparity coupled with parthenogenesis, producing clonal live offspring under favorable conditions (Bermingham and Wilkinson, 2009; Fouks et al., 2023; Simon et al., 2010). Certain earwigs (*Arixenia esau*) provide extended maternal care to their developing embryos internally (Tworzydlo et al., 2013; Tworzydlo et al., 2019), while some beetles and flies show morphological adaptations for nourishing intrauterine larvae, although the underlying mechanisms of this system are not well elucidated (Benoit et al., 2015; Ostrovsky et al., 2016).

By buffering embryos in a protected environment and sustained maternal investment, viviparity enhances offspring survival (Benoit et al., 2015; Dupoué et al., 2020; Lee and Harmer, 1980; Lodé, 2012; Ostrovsky et al., 2016) and allows species to exploit ecological niches that may be less hospitable to oviparous/ovoviviparous insects that lack prolonged pregnancy. Across vertebrates, the repeated evolution of viviparity has frequently been linked to environmental pressures, including low temperatures, variable climates, and other conditions that reduce the survival of externally developing eggs (Lambert and Wiens, 2013; Shine, 1995). More broadly, evolutionary theory predicts that increased parental investment should be favored when it improves offspring survival during vulnerable developmental stages, particularly where juvenile mortality is high (Benoit et al., 2019; Clutton-Brock, 1991). In insects, however, the ecological drivers of viviparity remain comparatively understudied, and experimental tests to elucidate these factors are rare. Compared to vertebrates, insects are especially susceptible to environmental stressors, such as desiccation, due to their small body size and the vulnerability of exposed eggs (Benoit et al., 2023). Retaining embryos within the maternal body may therefore protect against water loss, temperature fluctuations, predation, and other abiotic challenges, offering a potentially important adaptive advantage in harsh or unpredictable environments. Although these hypotheses have been suggested, few studies have directly tested improved survival and benefits underlying viviparity in insects.

Although viviparity has evolved repeatedly across animal lineages and is widely hypothesized to represent an adaptation to environmental stress, including drought, experimental studies directly testing the role of drought in the evolution and maintenance of viviparity remain scarce, particularly in insects. In this study, we compared viviparous *D. punctata* with two ovoviviparous cockroaches (the Madeira cockroach, *Rhyparobia maderae*, and the Lobster cockroach, *Nauphoeta cinerea*) in terms of potential survival during dry periods. The Pacific beetle-mimic cockroach *D. punctata* exhibits a highly derived reproductive mode that parallels aspects of mammalian pregnancy (Benoit et al., 2015; Fouks et al., 2023; Marchal et al., 2013). Unlike most cockroaches that are oviparous or ovoviviparous (produce eggs that develop and hatch inside the mother’s body, but receive no direct nutrients from the mother outside egg yolk), *D. punctata* is truly viviparous: females retain fertilized eggs within a specialized brood sac where embryos complete development (Marchal et al., 2013). During this time, the mother provides extensive provisioning to their developing embryos through a pseudoplacental structure and, most notably, by producing nutrient-rich “milk” secretions composed of proteins, lipids, and carbohydrates (Ingram et al., 1977; Stay and Coop, 1974; Williford et al., 2004). After midgut development, embryos consume these maternal secretions and package them into a crystalline form that serves as their sole food source (Banerjee et al., 2016; Santhakumari et al., 2023), enabling rapid growth and high survival rates. Once development is complete, the female gives birth to live, late-stage nymphs, which are less vulnerable than eggs or small, earlier instars. This strategy is among the most advanced reproductive systems in insects (Benoit et al., 2015; Fouks et al., 2023) and has made *D. punctata* a powerful model for investigating the evolution of viviparity.

*D. punctata* inhabits subtropical islands where drought conditions are frequent and often prolonged (Beard et al., 2005; Iese et al., 2021; McGree et al., 2016). These environments impose strong pressures that likely shape the cockroach’s biology and ecology, particularly regarding water conservation, reproductive investment, and survival strategies. On islands prone to periodic desiccation, *D. punctata* may benefit from its unique mode of reproduction, which allows offspring to bypass the vulnerable stages (eggs and early nymphal stages) and reduces risks associated with fluctuating water availability.

By adopting a comparative framework across three species, this study begins to address the gap while recognizing that the limited number of taxa necessitates cautious interpretation of broader evolutionary inferences. Based on our results, pregnant *D. punctata* females can support embryonic development for almost two weeks without access to water, and early nymphal stages are less susceptible to dehydration than those of ovoviviparous counterparts. These results indicate two potential advantages of viviparity: first, allowing embryos to survive for almost two weeks in dry habitats and second, bypassing early nymphal stages to reduce progeny susceptibility to dehydration. These factors may benefit viviparous species during rapid changes in water availability, suggesting that increased drought resistance could contribute to the shift to live birth under specific conditions and help prevent drought-induced declines in progeny survival.

## Materials and Methods

### Study species and husbandry

*R. maderae* and *N. cinerea* are useful comparisons to *D. punctata* because all three species retain developing eggs. However, unlike *D. punctata*, which nourishes embryos with milk-like secretions, the comparison species lack this specialized provisioning and give birth immediately after nymphs emerge from eggs, allowing study of traits specific to prolonged viviparity. Colonies of the Pacific Beetle mimic cockroach (*D. punctata*), the Madeira cockroach (*R. maderae*), and the Lobster cockroach (*N. cinerea*) were maintained at 25 °C under 70–80% relative humidity (RH) with a constant 12:12 h light:dark photoperiod. Colonies were provided with cardboard tubes for shelter and maintained on a consistent diet of Old Roy dog food and Tetramin fish food (Frigard et al., 2025; Jennings et al., 2019).

### Determination of water loss rates and survival among 3 cockroach species

We conducted a desiccation experiment in adults and juveniles of three species with different reproductive modes (two ovoviviparous, one viviparous) to compare mass change, water loss, and survival under desiccation. At 0-5% relative humidity (RH), changes in body mass were assumed to reflect only water loss because no water vapor could be absorbed from the surrounding environment (Benoit et al., 2005; Benoit et al., 2007). The low RH was maintained in a sealed chamber containing calcium sulfate (Drierite®) as a desiccant. The calcium sulfate continuously absorbed moisture from the air within the chamber, thereby maintaining a dry atmosphere. The RH inside the chamber was monitored using a hygrometer to be 0- 5% RH (listed as 0% RH hereafter).

Cockroaches were maintained at 0% RH and 25 °C, and individuals were weighed (Mettler Toledo® MS104TS/A00) at regular intervals (12 hours) throughout the exposure period to monitor progressive dehydration over four days. Following the experiment, dead individuals were dried to a constant mass at 90 °C under 0% RH to determine dry mass, and total body water content was calculated as the difference between fresh and dry mass. Rates of water loss, which include both cuticular transpiration and respiratory water loss, were estimated using the exponential model (Wharton, 1985). Specifically, the natural logarithm of the ratio of body mass at a given time to initial body mass was plotted against time, and the slope of the resulting regression line was used to estimate the water loss rate. Water loss rates were expressed as percent body water lost per hour (%/h), providing a standardized measure of transpiration under desiccating conditions. To assess survival during desiccation, individuals were placed in separate containers at 33% RH maintained in a desiccator with a saturated solution of MgCl_2_ (Winston and Bates, 1960). Survival was monitored and recorded at 12-hour intervals until all individuals had died or the observation ended.

### Progeny production and survival under periodic drought across species

To assess the effects of periodic dehydration on offspring survival, female ovoviviparous and viviparous cockroaches were assigned to either a hydrated control treatment or an intermittent dehydration treatment. Females were housed individually in plastic containers (11.6 × 10.2 × 3.8 cm; Pioneer Plastics), given a cardboard harborage, and maintained under standard colony conditions of 25 °C, 70–80% relative humidity, and a 12:12 h light:dark photoperiod. Control females had continuous access to food and water throughout gestation. Food consisted of dog food (Old Roy) supplied ad libitum, and water was provided using moistened cotton plugs replenished regularly. Females assigned to the dehydration treatment received continuous access to food but were exposed to repeated one-week periods without water. Specifically, water was removed for 7 consecutive days, then restored for a 7-day recovery period before the dehydration cycle was repeated. This intermittent dehydration regimen continued throughout reproduction for 100 days. Food remained continuously available throughout all treatment periods to ensure that observed effects were attributable specifically to water limitation rather than to nutritional deprivation. Progeny production was quantified by counting the total number of live nymphs produced at 10-day intervals for 100 days. At the conclusion of the experiment, the number of surviving progeny produced by hydrated and periodically dehydrated females was compared to determine whether repeated water limitation during pregnancy influences offspring production and viability in any of the three cockroach species.

### Osmolality assessment in *D. punctata* during pregnancy

Female *D. punctata* were randomly selected from laboratory colonies. The pregnancy stage was initially estimated from external morphological characteristics, including abdominal enlargement and the presence of gestation-associated white banding, and was subsequently verified either by birth/abortion or by dissection (Jennings et al., 2019; Jennings et al., 2020). Based on pregnancy status, females were randomly assigned to one of four treatment groups: early pregnancy under wet conditions, early pregnancy under dry conditions, late pregnancy under wet conditions, or late pregnancy under dry conditions. Females maintained under wet conditions were provided food and water ad libitum, whereas females subjected to dry conditions received food only and no access to water. Individuals were fed exclusively with dog food (Old Roy) throughout the experiments. Survival, as described earlier, was assessed in early- and late-pregnancy females.

Late-pregnancy females in wet and dry treatments were maintained under their respective conditions for 4, 8, 10, 12, and 14 days. Wet and dry treatments within each pregnancy stage were conducted concurrently to permit direct comparisons of dehydration responses. At the end of each treatment, females were anesthetized using CO2 and dissected on a non-porous surface. Embryo broods were separated from maternal tissues during dissection. The pregnancy stage was confirmed according to the previously described embryonic developmental criteria (Jennings et al., 2019). Embryo broods were transferred individually into 1.5 mL centrifuge tubes and stored at –80 °C until processing. Adult females were stored separately under identical conditions.

Hemolymph osmolality was measured to assess water balance and characterize physiological responses to dehydration. Adult cockroach samples were homogenized using a Tissue Tearor homogenizer (Biospec). Each sample was initially homogenized in 50 µL of deionized water. Embryo broods were homogenized separately using Disposable Pellet Pestles (Grainger) following the same procedure. Homogenized samples were transferred to the upper chamber of spin columns and centrifuged at 15,000 rpm for 25 min to isolate hemolymph. Extracted hemolymph samples were analyzed for osmolality using a vapor pressure osmometer (Wescor Vapro 5600) following the described methods (Benoit et al., 2014b; Rathore et al., 2024). For each sample, we obtained three independent osmolality readings and used the mean for subsequent analyses.

### Survival of *D. punctata* during early and late pregnancy

To assess the effects of drought and dehydration on survival during pregnancy, we conducted six independent survival trials: three in early pregnancy and three in late pregnancy. Female *D. punctata* were randomly isolated from colony populations and housed individually in experimental containers under controlled laboratory conditions. The pregnancy stage was initially determined using external morphological characteristics as described above.

For each trial, ten pregnant adult females were assigned to the dehydration treatment. Individuals subjected to dehydration stress were provided ad libitum access to dry dog food (Old Roy) as a nutritional source but denied free water throughout the experiment. This treatment was designed to simulate prolonged drought-like conditions and evaluate the capacity of pregnant females to tolerate water deprivation during different stages of gestation. Food was replenished as needed to ensure that mortality resulted primarily from dehydration stress rather than starvation.

Experimental containers were maintained under standardized environmental conditions consistent with colony maintenance, and all trials were conducted concurrently to minimize variation from environmental fluctuations. Survival was monitored daily by visually confirming the movement and responsiveness of each specimen. The number of living individuals remaining in each treatment group was recorded every 24 h until all specimens had died. These data were subsequently used to compare survival dynamics between early- and late-pregnant females under dehydrating conditions and to evaluate how gestational stage influences drought tolerance in viviparous cockroaches.

### Pregnancy length and progeny number in relation to mid-pregnancy drought

To establish if dehydration stress in mid-pregnancy affects gestational length and reproductive output, we exposed pregnant females to two experimental conditions: 1) dehydration stress for 8 days during mid-pregnancy, followed by ad libitum water, and 2) control females given ad libitum water throughout pregnancy. At the final juvenile instar, as females approached adult emergence, they were exposed to multiple adult males to maximize the likelihood of successful copulation and fertilization. Mating pairs were monitored regularly, and females that successfully mated were subsequently isolated for reproductive analyses. This approach ensured that pregnancy timing could be accurately tracked from fertilization through parturition. At 40 days of pregnancy, when milk production begins (Jennings et al., 2020; Stay and Coop, 1974), water was removed for 8 days, as previously described. Control individuals were allowed access to water throughout pregnancy. Recovery individuals had access to water for one week after dehydration, before sample collection for qPCR analysis of milk production.

Following successful mating, pregnant females were housed individually in plastic containers (11.6 × 10.2 × 3.8 cm; Pioneer Plastics) maintained under standard colony conditions as described above, including controlled temperature, humidity, and photoperiod. Each container included a small cardboard harborage to minimize stress and provide shelter during gestation. Females were provided ad libitum access to food and water throughout pregnancy (apart from the experimental period at 40 days). Individual housing prevented interactions among females and allowed accurate monitoring of reproductive timing and offspring production for each specimen.

Gestational length was defined as the number of days between observed mating and successful parturition. Females were monitored daily for signs of birth or abortion, and the timing of parturition was recorded daily upon the appearance of first-instar nymphs. Individuals that failed to produce offspring within 110 days after mating were excluded from subsequent analyses because these cases likely represented failed pregnancies or abortion of the initial brood. This exclusion threshold was selected based on previous reports indicating that normal pregnancy duration in this species rarely exceeds approximately 100 days (Engelmann, 1959), with most pregnancies occurring in 80-90 days. Establishing this cutoff reduced the likelihood of including abnormal reproductive events that could confound the interpretation of gestational timing. Reproductive output was quantified by counting the total number of live first-instar nymphs produced by each female immediately following birth. Only viable offspring were included in progeny counts. The final sample sizes for both the gestational length and progeny production analyses were 12 successfully reproducing females per treatment.

### Quantifying the effect of mid-pregnancy desiccation on milk protein transcription

To investigate the physiological mechanisms underlying pregnancy in *D. punctata*, we used qPCR to quantify expression of the known primary milk protein gene 13Y in a subset of samples from the previous section in which females were exposed to bouts of dehydration at 40 days of pregnancy. These proteins play critical roles in embryonic nourishment and have been implicated in regulating gestational timing in other viviparous insects (Benoit et al., 2012; Frigard et al., 2025; Jennings et al., 2020).

Mothers were collected at 48 days into pregnancy, a period corresponding to advanced embryonic development and active maternal nutrient provisioning. Samples were rapidly dissected to separate out embryos, flash frozen, and stored at −80 °C until molecular analyses were conducted. To minimize variation among individuals and obtain sufficient RNA yield, biological replicates were pooled from two pregnant females, with 6–8 independent biological replicates generated for each group.

Total RNA was extracted using TRIzol reagent (Invitrogen) according to the manufacturer’s recommended protocol. RNA quantity and quality were assessed using a NanoDrop before downstream analyses to ensure consistency among samples.

Complementary DNA (cDNA) synthesis was performed using 1 µg of total RNA with a cDNA Synthesis Kit (Thermo Scientific), generating templates for quantitative expression analysis. Quantitative PCR (qPCR) reactions were carried out using KiCqStart SYBR Green qPCR ReadyMix (Sigma-Aldrich) in combination with gene-specific primers targeting milk protein transcripts. Amplification reactions were performed on an Illumina Eco qPCR system under cycling conditions following previously established protocols (Frigard et al., 2025; Jennings et al., 2020).

Transcript abundance for the primary milk protein gene (NCBI accession: AY447988.1) was quantified for each biological replicate. Relative expression levels were normalized using the ΔΔCq method with the same validated reference genes and primer sets described previously (Jennings et al., 2020).

### Statistics and data analysis

All statistical analyses were performed in R 4.6.0 (Team, 2019). The Shapiro test (dplyr) for normality and the Breusch-Pagan and White tests (whitestrap) for heteroskedasticity were performed on each dataset. Normally distributed sets were analyzed using ANOVA (dplyr) and paired T-tests (dplyr), and non-normal sets using the Kruskal-Wallis test (dplyr) and Dunn’s test (FSA). For specific analyses, generalized linear models were utilized. Where possible, heteroskedasticity and abnormality were corrected using a log transformation. A Bonferroni correction was used when multiple groups were compared. R packages were utilized for data processing and visualization: ggplot2 (Wickham, 2016), viridis (Garnier et al., 2023), lmtest (Zeileis and Hothorn, 2015), whitestrap (Perez et al., 2020), dplyr (Wickham et al., 2023), FSA (Ogle et al., 2015), devtools (Wickham et al., 2022), and tidyverse (Wickham et al., 2019).

## Results

### Mass change, water loss, and survival differences among three cockroach species (two ovoviviparous and one viviparous)

Water balance traits differed significantly among the three cockroach species examined (Fig. 1). *D. punctata* exhibited the lowest proportional mass difference between 1st and 4th instars (d.f.=2,60; P < 0.05; Fig. 1A), suggesting that 1st instar *D. punctata* are relatively larger than the other two species (*N. cinerea* and *R. maderae*) during the first nymphal stage after birth. A higher body mass proportion in juvenile cockroaches for *D. punctata* relative to adults can improve water balance by reducing their surface area-to-volume ratio (Benoit and Denlinger, 2010; Wharton, 1985), thereby decreasing water loss through evaporation. This adaptation helps juveniles retain water more effectively, increasing their survival in dry or water-limited environments.

**Figure 1:**
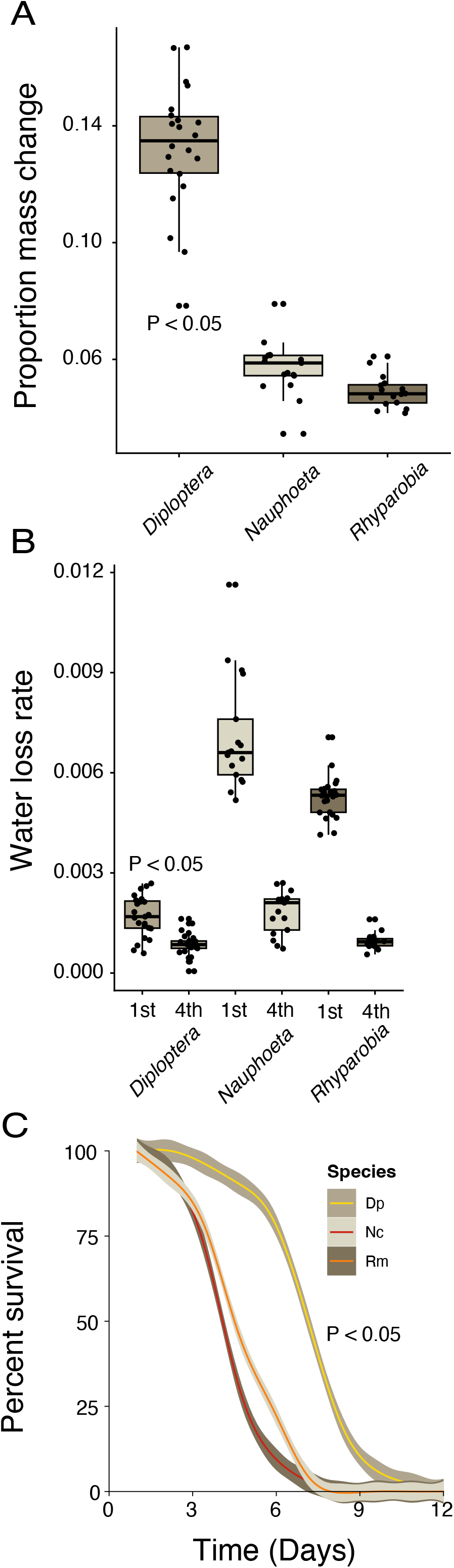
Water balance characteristics and desiccation survival of *Diploptera punctata*, *Nauphoeta cinerea*, and *Rhyparobia maderae*. (A) Proportional mass change for the three species in relation to developmental stage (first and fourth instars of each species). N = 20-22. (B) Water loss rates were measured in the first and fourth instars of each species. N = 20-24. (C) Survival curves under desiccating conditions for *D. punctata* (Dp), *N. cinerea* (Nc), and *R. maderae* (Rm) over 12 days. N = 3 of 20 cockroaches. Boxplots show medians, interquartile ranges, and individual biological replicates. Shaded regions in survival curves represent confidence intervals. Significant differences among treatments or species are indicated based on generalized linear models.

As expected, water loss rates varied significantly across developmental stages, with first instars consistently showing higher water loss than fourth instars (d.f.=2,32; P < 0.05; Fig. 1B) in all 3 species, but the extent of decline varied among species. Among the species tested, first instars of *N. cinerea* had the highest water loss rates. Survival of the first instars under desiccating conditions differed markedly among species, with *D. punctata* surviving substantially longer than both *N. cinerea* and *R. maderae* (d.f.=2,9; P < 0.05; Fig. 1C). Survival in *N. cinerea* and *R. maderae* declined rapidly after approximately 3–4 days, whereas *D. punctata* maintained high survival until approximately 6–7 days before declining. Overall, these studies indicate that juvenile *D. punctata* is more resistant to dehydration, most likely due to their larger size compared to *N. cinerea* and *R. maderae*.

### Progeny output and survival under simulated periods of drought across three cockroach species

Water availability strongly influenced progeny production across species, with dry conditions featuring extended periods that substantially reduced reproductive output in *N. cinerea* and *R. maderae*, but had comparatively limited effects on *D. punctata*. For *D. punctata*, progeny numbers increased steadily over time under both wet and dry conditions, with minor differences between treatments throughout the 100-day observation period (d.f.=1,4; P > 0.05, Fig. 2A). In contrast, *N. cinerea* and *R. maderae* exhibited pronounced reductions in accumulated progeny under dry conditions, whereas wet conditions supported rapid and sustained increases in offspring number over time at levels much higher than *D. punctata* (d.f.=1,4; P < 0.05, Fig. 2B,C). By the end of the experiment, wet-treated *N. cinerea* and *R. maderae* produced several-fold more progeny than dry-treated individuals. Comparisons of total progeny production further highlighted species-specific responses to dehydration, with *D. punctata* maintaining similar numbers of surviving across treatments (d.f.=2,6; P < 0.05, Fig. 2D), while *R. maderae* and especially *N. cinerea* experienced substantial declines under dry conditions (d.f.=2,6; P > 0.05, Fig. 2D). These results indicate that reproductive performance and progeny survival under dehydration vary markedly among species, with *D. punctata* exhibiting greater progeny output under dry environments.

**Figure 2:**
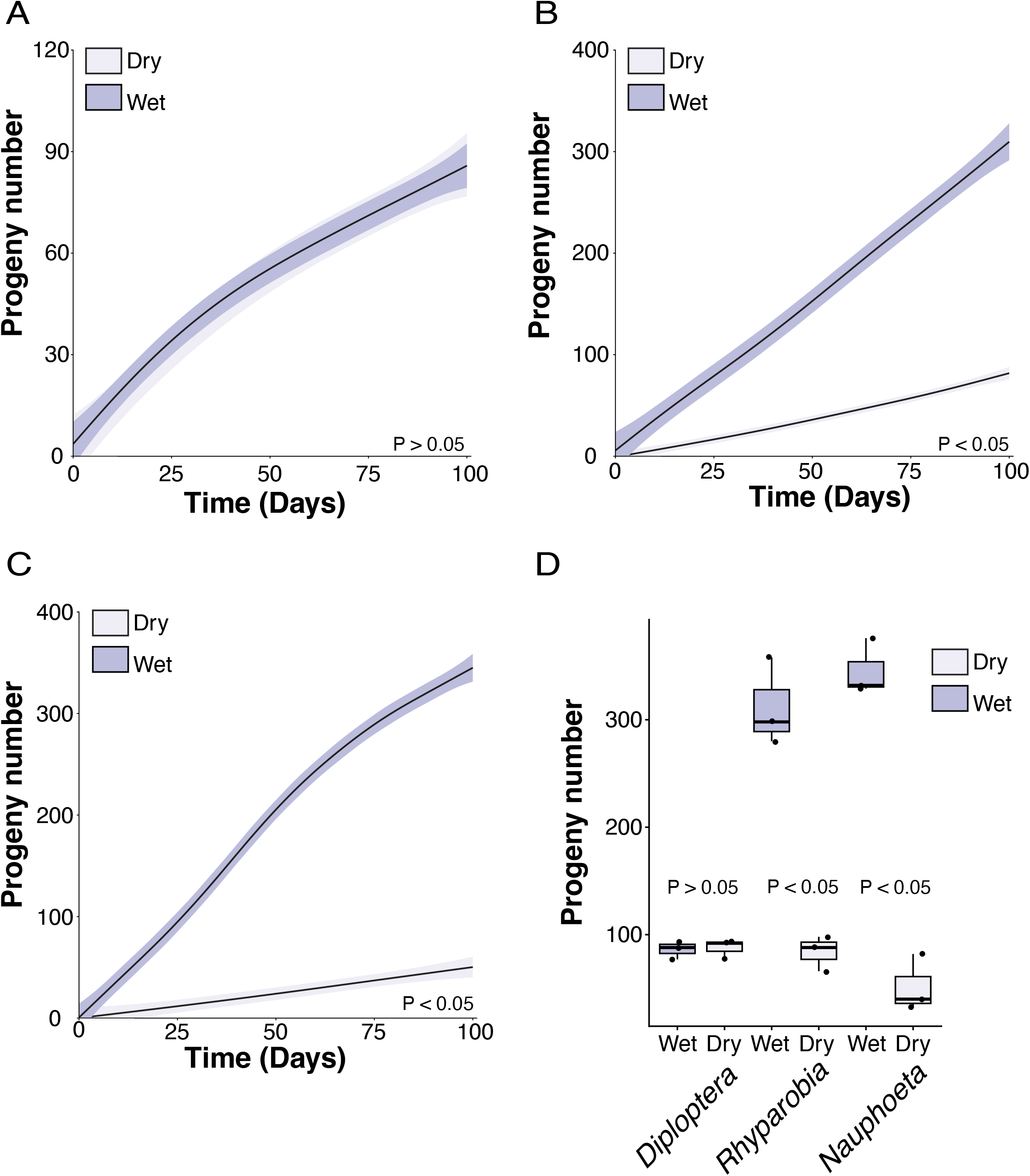
Effects of humidity conditions on progeny production in three cockroach species. (A) Cumulative progeny production over time in *Diploptera punctata* maintained under dry and wet conditions. N = 3 per wet and dry conditions. (B) Cumulative progeny production over time in *Rhyparobia maderae* under dry and wet conditions. N = 3 per wet and dry conditions. (C) Cumulative progeny production over time in *Nauphoeta cinerea* under dry and wet conditions. Shaded regions around fitted lines represent 95% confidence intervals. N = 3 per wet and dry conditions. (D) Total progeny production for each species under wet and dry conditions at the conclusion of the experiment. Boxplots indicate median values, interquartile ranges, and individual biological replicates. Significant differences among treatments or species are based on generalized linear models for three replicated dehydration experiments.

### *D. punctata* mothers buffer embryo osmolality during periods of desiccation stress

Prolonged dry conditions negatively affected survival, water balance, and reproductive success over time. Pregnant females (regardless of gestational stage) maintained under dry conditions had similar levels of survival (d.f.=1,4; P > 0.05, Fig. 3A). Hemolymph osmolality increased substantially in individuals exposed to dry conditions, particularly after 10–14 days, whereas osmolality remained relatively stable under wet conditions (d.f. = 6,81; P < 0.05, Fig. 3B).

**Figure 3:**
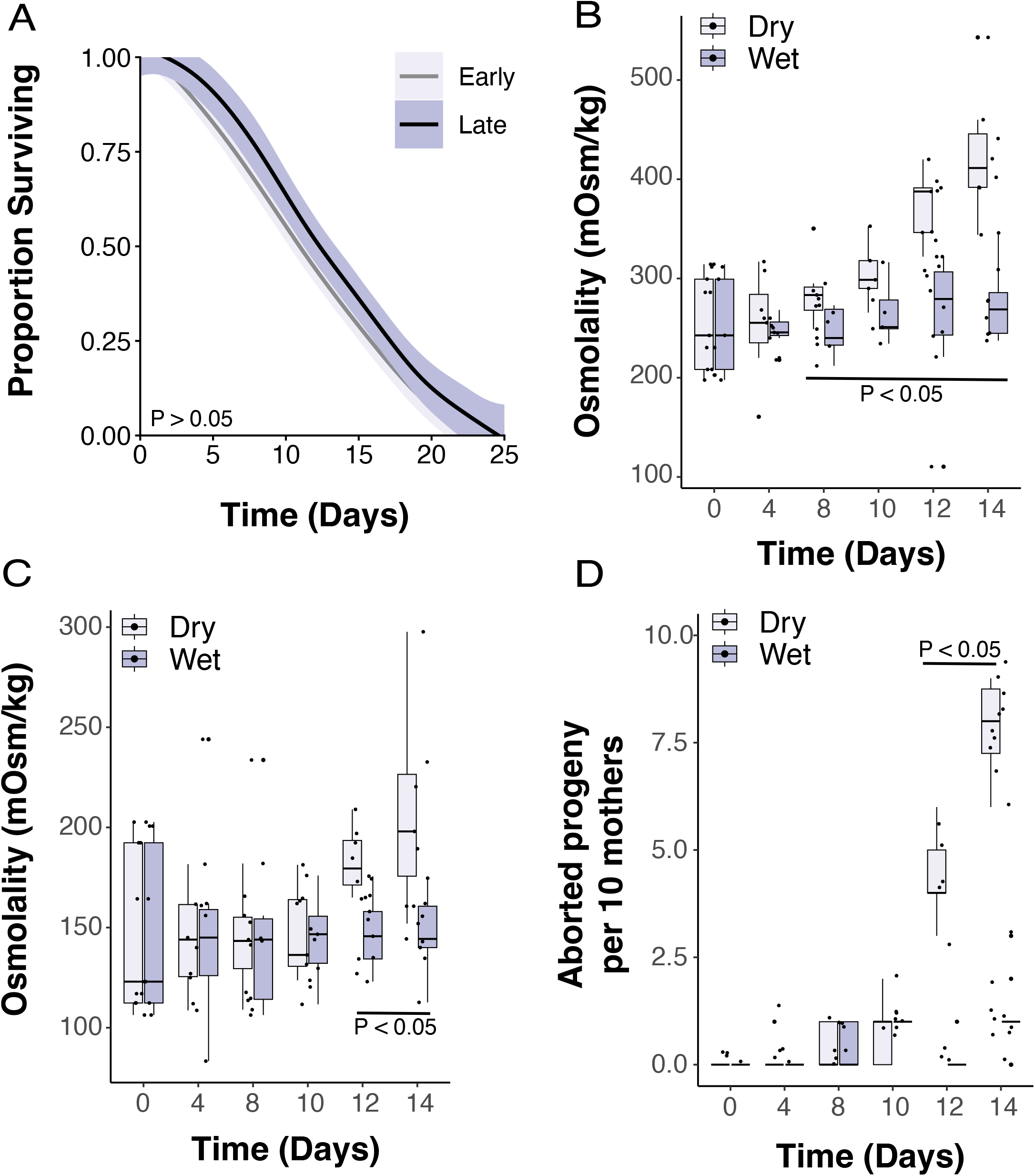
Effects of dehydration on survival, osmolality, and reproductive success of *Diploptera punctata.* (A) Survival of individuals maintained under early and late pregnancy over a 24-day period. N = 6 of 10 cockroaches. (B) Changes in maternal hemolymph osmolality under dry and wet conditions across time. (C) Progeny osmolality measured from offspring produced under dry and wet conditions over the course of the experiment. N= 8-10 replicates per treatment. (D) Number of aborted progeny per 10 mothers under dry and wet conditions across time. N = 5 of ten cockroaches. Boxplots indicate median values, interquartile ranges, and individual biological replicates. Shaded regions in survival curves represent confidence intervals around the fitted survival estimates. Significant differences among treatments or species are based on generalized linear models.

Similar trends were observed for embryo osmolality, with offspring from dry- exposed mothers exhibiting elevated osmolality values late in the exposure period (after ∼12 days) compared to embryos from mothers exposed to wet conditions, representing a control (d.f. = 6,85; P < 0.05, Fig. 3C). Reproductive success was strongly impaired by dehydration, as the number of aborted progeny increased progressively under dry conditions, reaching the highest levels (over 75% of females aborting) by day 14, while abortion rates remained minimal under wet conditions throughout the experiment (d.f. = 1,69; P < 0.05, Fig. 3D). Together, these results indicate that dehydration stress disrupts maternal water balance, whereas offspring are more buffered against water loss until extreme dehydration (10-12 days with no water) where the mother can no longer support the progeny and aborts.

### Adaptability in the *D. punctata* pregnancy cycle to accommodate dehydration stress

Dehydration exposure for 8 days altered reproductive timing and the expression of primary milk protein gene 13Y, but had limited effects on total progeny production.

Females exposed to dehydration exhibited a prolonged reproductive period, with the time to parturition increasing by 14% compared to the control (d.f.=1,21; P < 0.05, Fig. 4A). Despite this delay, the total number of progeny produced per female remained relatively similar between dehydrated and control groups, with overlapping distributions across treatments (d.f.=1,21; P > 0.05, Fig. 4B). Dehydration significantly reduced relative expression levels of the 13Y milk protein gene compared to controls, while recovery following dehydration partially restored expression toward control levels (d.f.=1,18; P < 0.05, Fig. 4C). These findings suggest that dehydration stress delays reproductive timing, possibly due to a reduced availability of milk as resources are diverted to female survival, but that some physiological effects are reversible following rehydration.

**Figure 4:**
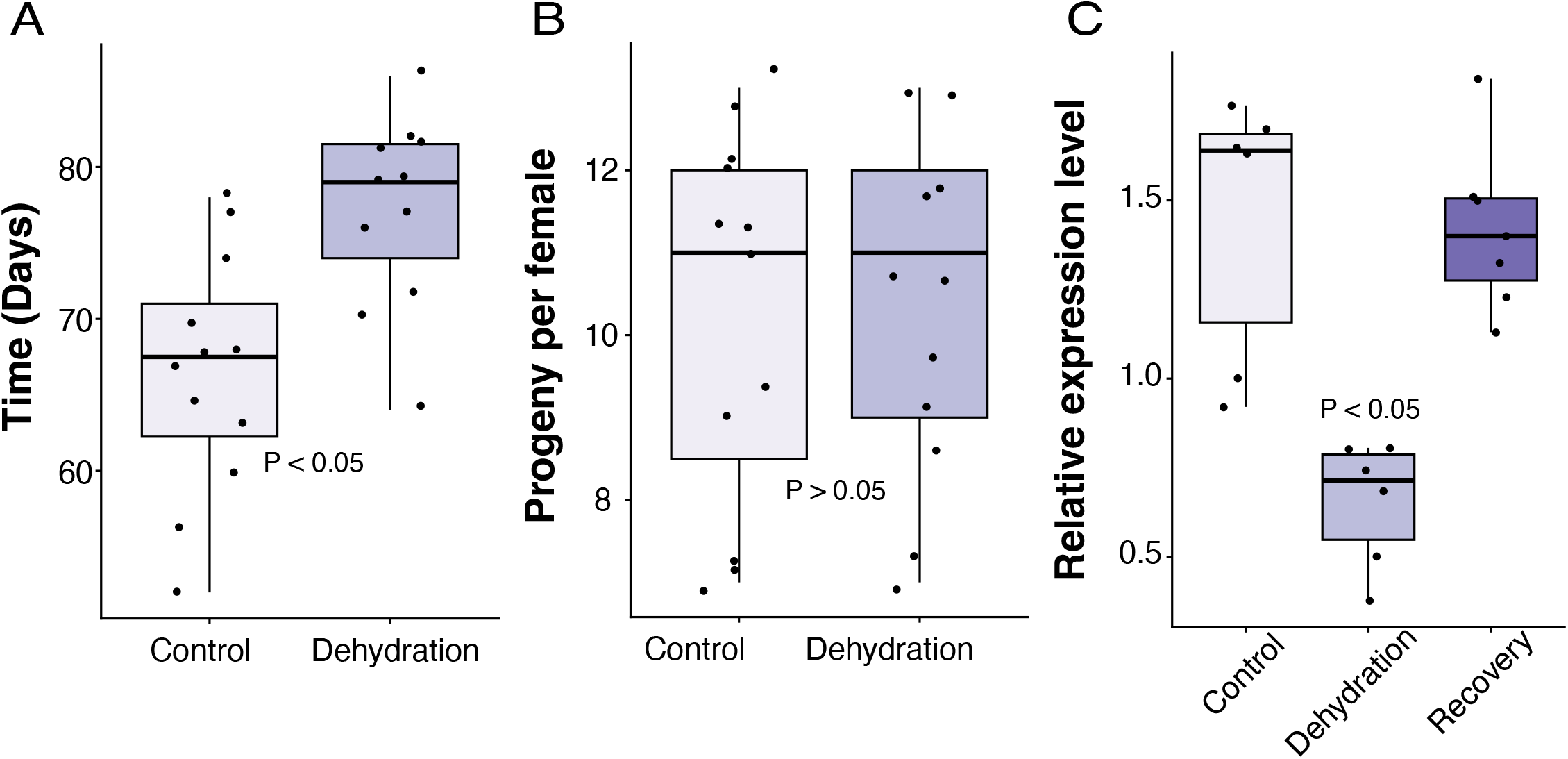
Effects of dehydration on reproductive timing, progeny production, and gene expression of *Diploptera punctata.* (A) Time to parturition for control and dehydrated individuals. N = 12 cockroaches. (B) Total progeny produced per female under control and dehydration treatments. N = 12 cockroaches. (C) Relative expression levels of the *milk protein* transcript under control, dehydration, and recovery conditions determined by quantitative PCR. N = 6 groups of two cockroaches. Boxplots indicate median values, interquartile ranges, and individual biological replicates. Recovery treatment groups represent individuals allowed to rehydrate following dehydration exposure. Significant differences among treatments or species are based on generalized linear models.

## Discussion

The repeated evolution of viviparity across insects and other animal taxa suggests that live birth offers significant ecological and physiological benefits under certain environmental conditions (Gavrilov-Zimin, 2022; Lodé, 2012). One proposed advantage is increased resistance to dehydration (Benoit et al., 2025; Lee and Harmer, 1980; Whittington et al., 2022), although experimental support for this hypothesis is lacking. In this study, *D. punctata* exhibited multiple traits consistent with the idea that viviparity buffers offspring against desiccation stress. Notably, pregnant females sustained embryonic development for nearly two weeks without access to water, maintaining stable embryonic osmolality to minimize dehydration-induced stress in their progeny.

These results indicate that the maternal environment protects developing intrauterine embryos from fluctuations in environmental water availability. Furthermore, *D. punctata* progeny are larger than those of ovoviviparous species and have significantly higher survival rates during dehydration stress immediately after birth. Together, these effects strongly support the idea that an extended pregnancy followed by live birth confers a distinct advantage for progeny during periods of low water availability. There may be some costs, such as delayed gestation or higher abortions, but these occur at points where it is likely juvenile stages outside the mother could have already succumbed to dehydration (Whittington et al., 2022).

A key finding of this study is that pregnancy imposed minimal additional dehydration burden on viviparous *D. punctata* females. Pregnant individuals survived water deprivation at rates comparable to non-pregnant/early pregnant females, despite simultaneously supporting the hydration and growth of their embryos. Although maternal osmolality increased steadily during dehydration, embryo osmolality remained relatively stable until water deprivation became prolonged. This pattern suggests that mothers prioritize water allocation to embryos, effectively shielding them from dehydration during critical developmental stages. Similar protection during viviparity has been observed in a viviparous lizard (Dupoué et al., 2015; Dupoué et al., 2018; Dupoué et al., 2020).

Such maternal buffering likely represents a major advantage of viviparity in drought-prone or unpredictable environments, as externally developing eggs would be directly exposed to desiccation (Benoit et al., 2025; Sulikowska-Drozd et al., 2019). By retaining embryos and early developmental stages internally, females can regulate the developmental environment and mitigate the risk of water loss to their progeny during dry periods. Indeed, this strategy may also relax seasonal constraints on female reproduction. By provisioning progeny with sufficient nutrients and water for development internally and using their own nutrient and hydration stores and larger biomass as a buffer, females can reproduce across a broader range of environmental conditions, reducing dependence on seasonal fluctuations in rainfall or plant phenology, which has been suggested in tsetse flies (Haines et al., 2020). This ecological flexibility may represent an additional selective advantage of viviparity.

Beyond protecting embryos during development, viviparity appears to produce offspring better equipped to survive after birth in dry conditions. Newly born nymphs of *D. punctata* are substantially larger than those of many ovoviviparous cockroach species, and this increased size likely contributes to improved dehydration resistance. This is due to the fact that they are born at a later developmental stage than if they were oviparous or ovoviviparous (Marchal et al., 2013). Larger body size reduces surface-area-to-volume ratios (Benoit et al., 2023; Wharton, 1985), thereby lowering water loss rates and enhancing survival under desiccating conditions. In contrast, oviparous and ovoviviparous species typically produce smaller hatchlings that must immediately cope with environmental stressors and are highly vulnerable to dehydration. Thus, the protective effects of viviparity extend beyond embryogenesis and into the early postnatal period. A limitation of this study is that the compared species were phylogenetically diverse and there are limited viviparous cockroach species (Evangelista et al., 2019; Evangelista et al., 2024; Thomas et al., 2020); as such, a more targeted and expansive survey across more cockroach species including the full spectrum from oviparous through to viviparous may be necessary to confirm these patterns.

Dehydration did not come without cost: dehydration led to a prolonged pregnancy, likely due to reduced milk production and nutrient provision to embryos. These extensions in pregnancy have previously been observed following a period of stress exposure, mainly sleep deprivation, due to lower milk protein expression in multiple viviparous systems (Benoit et al., 2014a; Frigard et al., 2025; Jennings et al., 2020). Although extended gestation may represent a cost, it may also function as an adaptive strategy that allows birth under more optimal conditions. By slowing embryonic development under water-limited conditions, females may avoid giving birth in environments unfavorable for offspring survival (Maag et al., 2023; Roubinov et al., 2021). Importantly, the number of offspring at birth was maintained despite delayed development, indicating that mothers prioritize offspring survival over developmental speed, a pattern similar to that observed in a viviparous lizard (Dupoué et al., 2018). It remains possible, however, that this reflects a trade-off in offspring size, as we observed reduced milk protein synthesis, although confirming this would require direct measurements of offspring size. This maintenance of progeny number under dehydration stress supports the role of maternal buffering during viviparity. Similar patterns have been documented in other viviparous systems (Dupoué et al., 2018), where maternal investment stabilizes offspring development under stressful conditions, suggesting that delayed development can help preserve offspring survival.

Despite the physiological demands associated with viviparity, there was little evidence of parent–offspring conflict in response to dehydration. Maternal survival remained unchanged during pregnancy despite water deprivation, and embryos maintained stable osmolality longer than their mothers, suggesting that maternal physiology effectively buffers developing offspring against dehydration. The primary indication of conflict emerges only under prolonged water stress, where increased abortion rates after two weeks may reflect a threshold beyond which maternal investment can no longer be sustained without significant expense to the mothers. Rather than indicating active maternal–offspring conflict, the extended gestation observed during dehydration is more consistent with a shared physiological constraint, likely resulting from reduced production of nutritive milk *in utero*. However, this prolonged gestation could still impose an indirect fitness cost on mothers by delaying subsequent reproduction, reducing future offspring production, impairing other physiological processes, or reducing lifespan (De Loof, 2011; Gilbert and Manica, 2010; Jasienska, 2009), potential trade-offs that were not assessed in this study. Future work examining lifetime reproductive output under repeated dehydration stress would help determine whether these maternal costs contribute to evolutionary conflict over resource allocation during pregnancy.

Overall, these findings strongly support the hypothesis that viviparity enhances drought resistance through maternal buffering during embryogenesis compared to ovoviviparous species, which confers greater tolerance to dehydration after birth.

Viviparous lizards exhibit several similar traits that may improve survival during dry periods (Dupoué et al., 2015; Dupoué et al., 2018; Dupoué et al., 2020), suggesting that these benefits could extend to vertebrate systems. In environments such as subtropical island habitats, where water availability can fluctuate dramatically, the ability to retain and sustain embryos internally likely confers a significant advantage, even if this is balanced against costs to females in terms of fecundity under stable conditions and the energetics of provisioning offspring. More broadly, this study provides direct evidence that live birth functions not only as a reproductive strategy to increase investment into progeny but also as a buffering mechanism that promotes offspring survival under harsh and variable conditions.

## Acknowledgments

Partial funding was provided by the National Institute of Allergy and Infectious Diseases of the National Institutes of Health under Award Number R01AI148551 and by the National Science Foundation under DEB1654417 (to J.B.B. for shared computational and reusable equipment resources). S.E. and A.J.v.R were supported by a UKRI Future Leaders Fellowship (MR/W007711/1, to S.E.) and the Wübben Stiftung Wissenschaft.

## References

Attardo, G. M., Abd-Alla, A. M. M., Acosta-Serrano, A., Allen, J. E., Bateta, R., Benoit, J. B., Bourtzis, K., Caers, J., Caljon, G., Christensen, M. B., et al. (2019). Comparative genomic analysis of six *Glossina genomes*, vectors of African trypanosomes. Genome Biol. 20, 187.

Banerjee, S., Coussens, N. P., Gallat, F.-X., Sathyanarayanan, N., Srikanth, J., Yagi, K. J., Gray, J. S. S., Tobe, S. S., Stay, B., Chavas, L. M. G., et al. (2016). Structure of a heterogeneous, glycosylated, lipid-bound, in vivo-grown protein crystal at atomic resolution from the viviparous cockroach *Diploptera punctata*. IUCrJ 3, 282–293.

Beard, K. H., Vogt, K. A., Vogt, D. J., Scatena, F. N., Covich, A. P., Sigurdardottir, R., Siccama, T. G. and Crowl, T. A. (2005). Structural and functional responses of a subtropical forest to 10 years of hurricanes and droughts. Ecol. Monogr. 75, 345– 361.

Benoit, J. B. and Denlinger, D. L. (2010). Meeting the challenges of on-host and off-host water balance in blood-feeding arthropods. J. Insect Physiol. 56, 1366–1376.

Benoit, J. B., Yoder, J. A., Rellinger, E. J., Ark, J. T. and Keeney, G. D. (2005). Prolonged maintenance of water balance by adult females of the American spider beetle, *Mezium affine* Boieldieu, in the absence of food and water resources. J. Insect Physiol. 51, 565–573.

Benoit, J. B., Lopez-Martinez, G., Robert Michaud, M., Elnitsky, M. A., Lee, R. E. and Denlinger, D. L. (2007). Mechanisms to reduce dehydration stress in larvae of the Antarctic midge, *Belgica antarctica*. J. Insect Physiol. 53, 656–667.

Benoit, J. B., Attardo, G. M., Michalkova, V., Takác, P., Bohova, J. and Aksoy, S. (2012). Sphingomyelinase activity in mother’s milk is essential for juvenile development: a case from lactating tsetse flies. Biol. Reprod. 87, 17, 1–10.

Benoit, J. B., Attardo, G. M., Michalkova, V., Krause, T. B., Bohova, J., Zhang, Q., Baumann, A. A., Mireji, P. O., Takáč, P., Denlinger, D. L., et al. (2014a). A novel highly divergent protein family identified from a viviparous insect by RNA-seq analysis: a potential target for tsetse fly-specific abortifacients. PLoS Genet. 10, e1003874.

Benoit, J. B., Hansen, I. A., Attardo, G. M., Michalková, V., Mireji, P. O., Bargul, J. L., Drake, L. L., Masiga, D. K. and Aksoy, S. (2014b). Aquaporins are critical for provision of water during lactation and intrauterine progeny hydration to maintain tsetse fly reproductive success. PLoS Negl. Trop. Dis. 8, e2517.

Benoit, J. B., Attardo, G. M., Baumann, A. A., Michalkova, V. and Aksoy, S. (2015). Adenotrophic viviparity in tsetse flies: potential for population control and as an insect model for lactation. Annual Review of Entomology 60, 351–371.

Benoit, J. B., Kölliker, M. and Attardo, G. M. (2019). Putting invertebrate lactation in context. Science 363, 593.

Benoit, J. B., McCluney, K. E., DeGennaro, M. J. and Dow, J. A. T. (2023). Dehydration dynamics in terrestrial arthropods: from water sensing to trophic interactions. Annu. Rev. Entomol. 68, 129–149.

Benoit, J. B., Weaving, H., McLellan, C., Terblanche, J. S., Attardo, G. M. and English, S. (2025). Viviparity and obligate blood feeding: tsetse flies as a unique research system to study climate change. Curr. Opin. Insect Sci. 69, 101369.

Bermingham, J. and Wilkinson, T. L. (2009). Embryo nutrition in parthenogenetic viviparous aphids. Physiol. Entomol. 34, 103–109.

Clutton-Brock, T. H. (1991). The evolution of parental care. Princeton, NJ: Princeton University Press.

De Loof, A. (2011). Longevity and aging in insects: Is reproduction costly; cheap; beneficial or irrelevant? A critical evaluation of the “trade-off” concept. J. Insect Physiol. 57, 1–11.

Dupoué, A., Brischoux, F., Angelier, F., DeNardo, D. F., Wright, C. D., Lourdais, O. and Jennifer, G. (2015). Intergenerational trade-off for water may induce a mother– offspring conflict in favour of embryos in a viviparous snake. Funct. Ecol. 29, 414– 422.

Dupoué, A., Le Galliard, J.-F., Josserand, R., DeNardo, D. F., Decencière, B., Agostini, S., Haussy, C. and Meylan, S. (2018). Water restriction causes an intergenerational trade-off and delayed mother–offspring conflict in a viviparous lizard. Funct. Ecol. 32, 676–686.

Dupoué, A., Sorlin, M., Richard, M., Le Galliard, J. F., Lourdais, O., Clobert, J. and Aubret, F. (2020). Mother-offspring conflict for water and its mitigation in the oviparous form of the reproductively bimodal lizard, *Zootoca vivipara*. Biol. J. Linn. Soc. Lond. 129, 888–900.

Evangelista, D. A., Wipfler, B., Béthoux, O., Donath, A., Fujita, M., Kohli, M. K., Legendre, F., Liu, S., Machida, R., Misof, B., et al. (2019). An integrative phylogenomic approach illuminates the evolutionary history of cockroaches and termites (Blattodea). Proc. Biol. Sci. 286, 20182076.

Evangelista, D. A., Nelson, D., Kotyková Varadínová, Z., Kotyk, M., Rousseaux, N., Shanahan, T., Grandcolas, P. and Legendre, F. (2024). Phylogenomic analyses of Blattodea combining traditional methods, incremental tree-building, and quality-aware support. Mol. Phylogenet. Evol. 200, 108177.

Fouks, B., Harrison, M. C., Mikhailova, A. A., Marchal, E., English, S., Carruthers, M., Jennings, E. C., Chiamaka, E. L., Frigard, R. A., Pippel, M., et al. (2023). Live-bearing cockroach genome reveals convergent evolutionary mechanisms linked to viviparity in insects and beyond. iScience 26, 107832.

Frigard, R., Ajayi, O. M., LeFevre, G., Ezemuoka, L. C., English, S. and Benoit, J. B. (2025). Daily activity rhythms, sleep and pregnancy are fundamentally related in the Pacific beetle mimic cockroach, *Diploptera punctata*. J. Exp. Biol. 228,.

Garnier, S., Ross, N., Rudis, B., Filipovic-Pierucci, A., Galili, T., timelyportfolio, O’Callaghan, A., Greenwell, B., Sievert, C., Harris, D. J., et al. (2023). sjmgarnier/viridis: CRAN release v0.6.3. Zenodo.

Gavrilov-Zimin, I. A. (2022). Development of theoretical views on viviparity. Biol. Bull. Rev. 12, 570–595.

Gilbert, J. D. J. and Manica, A. (2010). Parental care trade-offs and life-history relationships in insects. Am. Nat. 176, 212–226.

Haines, L. R., Vale, G. A., Barreaux, A. M. G., Ellstrand, N. C., Hargrove, J. W. and English, S. (2020). Big baby, little mother: Tsetse flies are exceptions to the juvenile small size principle. Bioessays 42, e2000049.

Iese, V., Kiem, A. S., Mariner, A., Malsale, P., Tofaeono, T., Kirono, D. G. C., Round, V., Heady, C., Tigona, R., Veisa, F., et al. (2021). Historical and future drought impacts in the Pacific islands and atolls. Clim. Change 166,.

Ingram, M. J., Stay, B. and Cain, G. D. (1977). Composition of milk from the viviparous cockroach, Diploptera punctata. Insect Biochem. 7, 257–267.

Jasienska, G. (2009). Reproduction and lifespan: Trade-offs, overall energy budgets, intergenerational costs, and costs neglected by research. Am. J. Hum. Biol. 21, 524–532.

Jennings, E. C., Korthauer, M. W., Hamilton, T. L. and Benoit, J. B. (2019). Matrotrophic viviparity constrains microbiome acquisition during gestation in a live-bearing cockroach, *Diploptera punctata*. Ecol. Evol. 9, 10601–10614.

Jennings, E. C., Korthauer, M. W., Hendershot, J. M., Bailey, S. T., Weirauch, M. T., Ribeiro, J. M. C. and Benoit, J. B. (2020). Molecular mechanisms underlying milk production and viviparity in the cockroach, Diploptera punctata. Insect Biochem. Mol. Biol. 120, 103333.

Kalinka, A. T. (2015). How did viviparity originate and evolve? Of conflict, co-option, and cryptic choice. Bioessays 37, 721–731.

Lambert, S. M. and Wiens, J. J. (2013). Evolution of viviparity: a phylogenetic test of the cold-climate hypothesis in phrynosomatid lizards: Evolution of viviparity. Evolution 67, 2614–2630.

Lee, J. A. and Harmer, R. (1980). Vivipary, a reproductive strategy in response to environmental stress? Oikos 35, 254.

Lodé, T. (2012). Oviparity or viviparity? That is the question…. Reprod. Biol. 12, 259– 264.

Maag, N., Cozzi, G., Seager, D., Manser, M., Sickmüller, A., Hildebrandt, T. B., Clutton-Brock, T. and Ozgul, A. (2023). Dispersal-induced social stress prolongs gestation in wild meerkats. Biol. Lett. 19, 20230183.

Marchal, E., Hult, E. F., Huang, J., Stay, B. and Tobe, S. S. (2013). Diploptera punctata as a model for studying the endocrinology of arthropod reproduction and development. Gen. Comp. Endocrinol. 188, 85–93.

McGree, S., Schreider, S. and Kuleshov, Y. (2016). Trends and variability in droughts in the Pacific Islands and northeast Australia. J. Clim. 29, 8377–8397.

Ogle, D. H., Doll, J. C., Wheeler, A. P. and Dinno, A. (2015). FSA: Simple Fisheries Stock Assessment Methods.

Ostrovsky, A. N., Lidgard, S., Gordon, D. P., Schwaha, T., Genikhovich, G. and Ereskovsky, A. V. (2016). Matrotrophy and placentation in invertebrates: a new paradigm: Invertebrate matrotrophy and placentation. Biol. Rev. Camb. Philos. Soc. 91, 673–711.

Perez, J., Lopez, J. and Jeong, M. J. L. (2020). Package ‘whitestrap. *Package ‘whitestrap*.

Rathore, S., Mitra, A. T., Hyland-Brown, R., Jester, A., Layne, J. E., Benoit, J. B. and Buschbeck, E. K. (2024). Osmosis as nature’s method for establishing optical alignment. Curr. Biol. 34, 1569–1575.e3.

Roubinov, D., Meaney, M. J. and Boyce, W. T. (2021). Change of pace: How developmental tempo varies to accommodate failed provision of early needs. Neurosci. Biobehav. Rev. 131, 120–134.

Santhakumari, P. R., Dhanabalan, K., Virani, S., Hopf-Jannasch, A. S., Benoit, J. B., Chopra, G. and Subramanian, R. (2023). Variability in phenylalanine side chain conformations facilitates broad substrate tolerance of fatty acid binding in cockroach milk proteins. PLoS One 18, e0280009.

Shine, R. (1995). A new hypothesis for the evolution of viviparity in reptiles. Am. Nat. 145, 809–823.

Simon, J.-C., Stoeckel, S. and Tagu, D. (2010). Evolutionary and functional insights into reproductive strategies of aphids. C. R. Biol. 333, 488–496.

Stay, B. and Coop, A. C. (1974). “Milk” secretion for embryogenesis in a viviparous cockroach. Tissue Cell 6, 669–693.

Sulikowska-Drozd, A., Apostolopoulou, K., Giokas, S. and Schilthuizen, M. (2019). Viviparous reproduction in the land snail Idyla (Pulmonata: Clausiliidae) from Greece: a disadvantageous inheritance? J. Molluscan Stud. 85, 262–270.

Team, R. C. (2019). R: a language and environment for statistical computing, version 3.0. 2. Vienna, Austria: R Foundation for Statistical Computing; 2013.

Thomas, G. W. C., Dohmen, E., Hughes, D. S. T., Murali, S. C., Poelchau, M., Glastad, K., Anstead, C. A., Ayoub, N. A., Batterham, P., Bellair, M., et al. (2020). Gene content evolution in the arthropods. Genome Biol. 21, 1–14.

Tworzydlo, W., Kisiel, E. and Bilinski, S. M. (2013). Embryos of the viviparous dermapteran, Arixenia esau develop sequentially in two compartments: terminal ovarian follicles and the uterus. PLoS One 8, e64087.

Tworzydlo, W., Jaglarz, M. K., Pardyak, L., Bilinska, B. and Bilinski, S. M. (2019). Evolutionary origin and functioning of pregenital abdominal outgrowths in a viviparous insect, Arixenia esau. Sci. Rep. 9, 16090.

Wharton, G. W. (1985). Water Balance of Insects. *Regulation: Digestion, Nutrition*, Excretion 565–601.

Whittington, C. M., Van Dyke, J. U., Liang, S. Q. T., Edwards, S. V., Shine, R., Thompson, M. B. and Grueber, C. E. (2022). Understanding the evolution of viviparity using intraspecific variation in reproductive mode and transitional forms of pregnancy. Biol. Rev. Camb. Philos. Soc. 97, 1179–1192.

Wickham, H. (2016). Ggplot2. 2nd ed. Basel, Switzerland: Springer International Publishing.

Wickham, H., Averick, M., Bryan, J., Chang, W., McGowan, L., François, R., Grolemund, G., Hayes, A., Henry, L., Hester, J., et al. (2019). Welcome to the tidyverse. J. Open Source Softw. 4, 1686.

Wickham, H., Hester, J., Chang, W. and Bryan, J. (2022). devtools: Tools to make developing R packages easier.

Wickham, H., Fracois, R., Henry, L., Muller, K. and Vaughan, D. (2023). dplyr: A Grammar of Data Manipulation.

Williford, A., Stay, B. and Bhattacharya, D. (2004). Evolution of a novel function: nutritive milk in the viviparous cockroach, *Diploptera punctata*. Evol. Dev. 6, 67–77.

Winston, P. W. and Bates, D. H. (1960). Saturated solutions for the control of humidity in biological research. Ecology 41, 232–237.

Zeileis, A. and Hothorn, T. (2015). Diagnostic checking in regression relationships.

